# Bivariate confidence probability plots as a method to test the accuracy and variability of microbiological measures

**DOI:** 10.1101/2020.11.18.388603

**Authors:** Antonio Monleon-Getino, AC Marca Home Care

## Abstract

**Introduction:** In an interlaboratory calibration analysis to validate a methodology that will be proposed as a European standard for domestic laundry disinfection, tests were carried out to detect if there are different behaviors in the measurements regarding accuracies and variabilities. Interlaboratory tests using different doses of disinfectant and microorganisms were carried out. ISO 5725-2 and ISO 13528 form the basis of validations of quantitative methods, providing validation specifications for interlaboratory studies. However, a need for a simple graphical method to detect interlaboratory differences in accuracy and variability was observed.

**Objectives:** The general goal of this work is to present a new exploratory methodology, graphical and easy to interpret, that can determine the accuracy and variability (precision) of a variable, and compare it to the methodology applied in ISO 5725-2 and ISO 13528.

**Methods:** We used confidence probability plots of the multivariate Student’s t-distribution to observe the accuracy and variability of microbiological measures carried out by different laboratories during a ring trial exercise. A function in R was built for this purpose: Miriam.analysis.ellipse(Y, factor_a, eel.plot = “ t-Student”). The different observations of accuracy and variability are represented in the ellipses. If any of the points are outside the ellipse with 95% confidence, we can assume a deviation in accuracy and / or variability.

**Results:** Two examples are provided with real microbiological data (logarithmic unit reductions (LR) for *Pseudomonas aeruginosa, Escherichia coli, Staphilococcus aureus, Enterococcus hirae, Candida albicans* and microbial counts in water (WW)). The proposed new method allowed us to detect possible deviations in the WWMEA variable and we believe it has future application for the rapid control of microbiological measures.

## 1. Background

Microbiological systems have a high number of uncontrollable variables, which increases experimental complexity and reduces accuracy. The considerable number of variables at play could lead to misinterpretation of data or uncorrectable errors. The sources of variability in microbiology methods fall roughly into three groups: the test system (e.g. microorganism and environmental conditions), the scientist(s) performing the study, and in the case of antimicrobial efficacy studies, the test substance (1). Although various causes of variability have been defined, there may be others that are equally substantial but are unknown to microbiologists (1). In general, experimental results must be reproducible to be meaningful. As it is known that this variability exists in biological systems and that it is impossible to avoid, strategies should be sought to reduce its impact on significance.

A ring trial carried out to test the validity of a method to be proposed as a European standard for domestic laundry disinfection in terms of accuracy and variability was used to apply the proposed statistical approach.

During the ring trial, microbial reductions of a known population exposed to the product during a washing cycle were evaluated. Microbiological variables such as LR (logarithm reduction log (CFU/ml) for different types of microorganisms), WW (counts of microorganisms in the wash water) or RI (Recovery of microorganisms in the cross-contamination carrier) were generated and compared statistically between laboratories, laboratory-scale devices, product type and product dose. Accuracy and variability were defined for each variable

Below are the common definitions of accuracy and variability [4][5][6][7]. Accuracy has two definitions [4]:

- Most commonly, it is a description of systematic errors and a measure of statistical bias; low accuracy, referred to by ISO as trueness, generates difference between a result and a “true” value [5].
- Alternatively, ISO defines accuracy as describing a combination of both types of observational error (random and systematic), so high accuracy requires both high precision and high trueness.

Precision is a description of random errors and a measure of statistical variability [4].

## 2. Objectives

The goal of this work is to present a new exploratory methodology, graphical and easy to interpret, for observing the accuracy and variability (precision) of a variable.

## 3. Material and methods

We propose using confidence probability plots to test the accuracy and variability of microbiological measures made by different laboratories during a ring trial. The proposed plot represents a multivariate Student’s t-distribution for the bivariate distribution of the accuracy and variability of the measured variables (e.g. LR) for the different laboratories and a product dose.

Thus, the proposed method is based on a bivariate distribution, representing the coordinates, accuracy and variability for each observation and laboratory obtained during the ring trial. Afterwards, the bivariate confidence interval bands are estimated assuming a multivariate Student’s t-distribution and it is observed if any sample falls outside the 95% interval (agreed as acceptability of accuracy or variability)

The following is a summary of this distribution based on the summary in https://www.statlect.com/probability-distributions/multivariate-student-t-distribution

### Multivariate Student’s t-distribution

Student’s t-distribution is a continuous probability distribution that arises when estimating the mean of a normally distributed population (N(*μ,σ*)) in situations where the sample size is small (n<30) and the population standard deviation is unknown [9][10].

Let X={X_1_, …, X_n_} a vector of *n* observations of a target variable (e.g. LR for each microorganism), and *X* be independently and identically drawn from the distribution *X*~N(*μ,σ*). We assume this is a sample of size *n* from a normally distributed population with the expected mean value *μ* and unknown variance *σ*^2^.

t-distribution has the probability density function given by

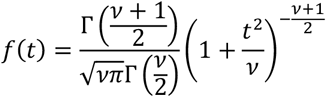

where v is the number of degrees of freedom and Γ is the gamma function. The overall shape of *f*(*t*) resembles the bell shape of a normally distributed variable with N(*μ*=*0,σ*=1), except that it is lower and wider. Student’s t-distribution leads to a variety of statistical estimation problems, where the goal is to estimate an unknown parameter, such as a mean value, in a setting where the data are observed with additive errors [9][10].

The multivariate Student’s t-distribution:

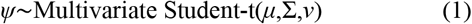

where μ is a vector of means, ∑ is a matrix of variances-covariances and *v* is the vector of degrees of freedom. It is a generalization of random vectors of the Student’s t-distribution, which is applicable to univariate random variables [11][12][13].

If a random variable, X, has a standard univariate Student’s t-distribution, *X* ~ t-distribution(*v*), it can be represented as a ratio between a standard normal random variable and the square root of a Gamma random variable. In the same way, a random vector has a standard multivariate Student’s t-distribution if it can be represented as a ratio between a standard multivariate normal random vector and the square root of a Gamma random variable [11] [12].

The procedure was encapsulated in the function for R [14]:

Miriam.analysis.ellipse(Y, factor_a, eel.plot = “ t-Student”)

Arguments:

- Y=response variable
- factor_a= vector with levels of factor where variability-accuracy between levels is analyzed
- eel.plot=type of ellipse eel.plot=“MV Normal”, or “MV t-Student”

Miriam.analysis.ellipse() uses the function stat_ellipse () with the argument type=“t” to compute a ellipse plot with bands representing confidence bands (5, 25, 50, 75, 95 and 99% of confidence) and assumes a multivariate t-distribution [15].

Miriam.analysis.ellipse() is a function of the R library called Diagnobatch() developed by Monleón (2020) [14]. Some of these methods are novel in microbiology but they have been widely used in other industrial sectors, such as the pharmaceutical industry, which, as occurs in this situation, presents a large number of redundant variables and where accuracy, variability and outliers hypotheses must be verified (using the common statistical methods: Cochran’s test, Fligner-Killeen test, Grubs test) in order to detect inconsistencies between groups (in this case laboratories) but using a multivariate approach and summarizing the variables through principal components analysis [16] [17].

## 4. Results

### Accuracy versus variability plots with confidence bands for each variable studied

The means (accuracy) and standard deviations (variability) of the values used by the laboratories have been represented in a plane, and a multivariate t-distribution has been superimposed using a ellipse plot. Below are some examples with real laboratory results obtained during the ring trial.

In Figures 1 and 2, an ellipse plot constructed with the function Miriam.analysis.ellipse() is represented for the results of variables LR for *P. aeruginosa* and WWMEA. Level lines represent a region of confidence between 5 and 99%. The common interval 95% was deemed to detect problems with accuracy or variability in the measures.

**Figure 1:**
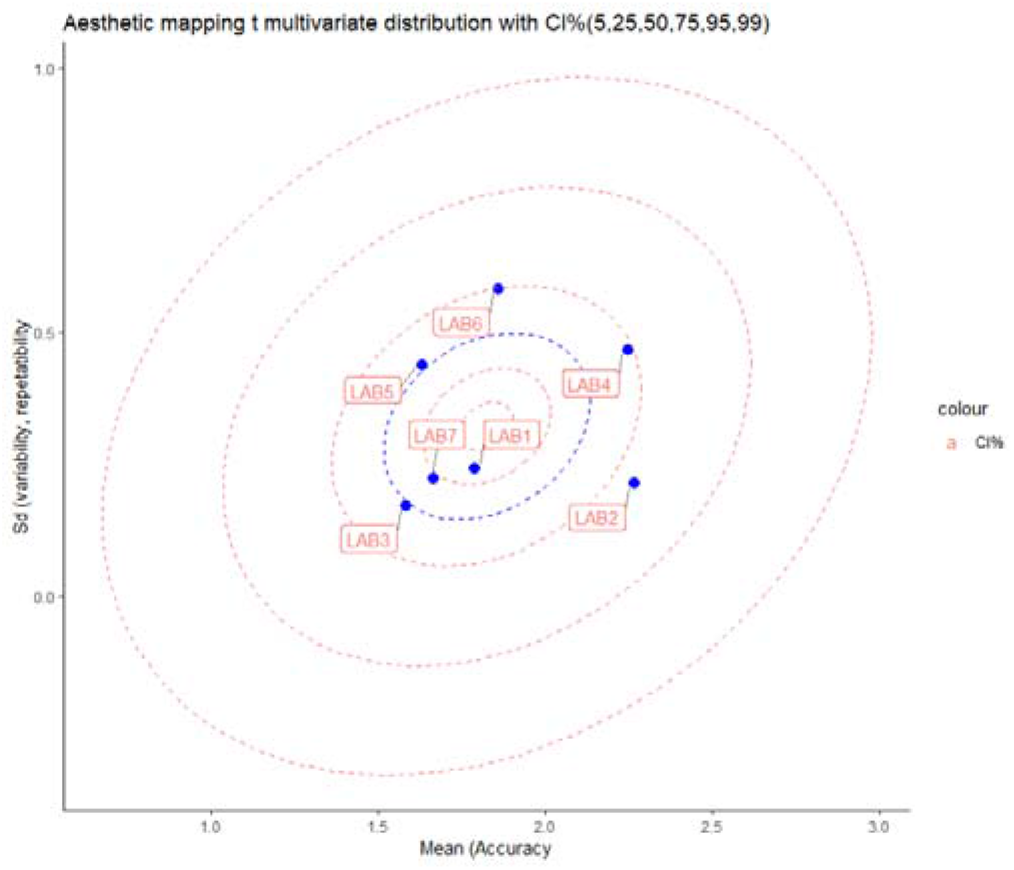
Accuracy versus variability plots with confidence bands supposing a multivariate Student’s t-distribution for the LR values obtained for *P. aeruginosa* in different samples of different laboratories and concentrations of disinfectant during a ring trial.

**Figure 2:**
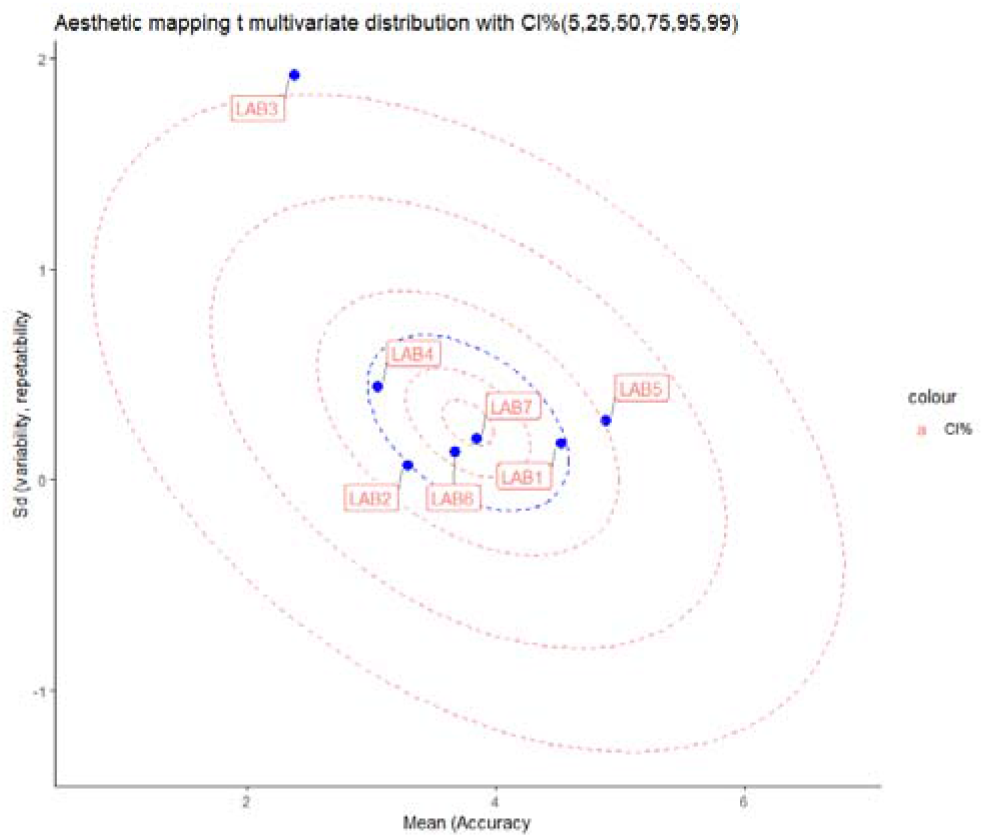
Accuracy versus variability plots with confidence bands supposing a multivariate Student’s t distribution for the values obtained for WWMEA in different samples of different laboratories and concentrations of disinfectant during a ring trial.

In Figure 1 it is possible to observe that all laboratory means and standard deviations for the LR of *P. aeuriginosa* are inside the confidence bands of 95%. It can therefore be concluded that in all cases variability and accuracy are in the same range of variation (ellipse center) with a confidence of 95%.

In the second case (Figure 2), high variability-accuracy (out of the 95%CI band) of laboratory 3 is detected for the variable WWMEA, providing evidence of differential behaviour from the other laboratories. In a later step, it would be necessary to investigate the measurements from this laboratory and perhaps repeat them to achieve compatibility with the other laboratories.

## 5. Conclusions

We believe that the proposed exploratory method constitutes a quick and graphic first step to observe differences between levels of a factor (e.g. laboratory accuracy and variability) and differential behaviors between levels.

The new function for R, Miriam.analysis.ellipse(Y, factor_a, eel.plot = “t-Student”), is proposed in order to compute a ellipse plot with bands representing confidence bands (5, 25, 50, 75, 95 and 99% of confidence) assuming a multivariate t-distribution.

In the proposed method, using Miriam.analysis.ellipse(Y, factor_a, eel.plot = “ t-Student”), the different observations of accuracy and variability are represented in the ellipses. If any of the points are outside the 95% confidence band, we can assume there is a problem with accuracy and / or variability.

Two examples are presented with real microbiological data (LR, WW) obtained during an interlaboratory calibration test with highly variable microbiological variables. The new method allowed us to detect problems with the WWMEA variable and we believe it has future application in controling microbiological measures.

